# Oxytocin increases after affiliative interactions in male Barbary macaques

**DOI:** 10.1101/695064

**Authors:** Alan V. Rincon, Tobias Deschner, Oliver Schülke, Julia Ostner

## Abstract

Mammals living in stable social groups often mitigate the costs of group living through the formation of social bonds and cooperative relationships. The neuropeptide hormone oxytocin (OT) has been proposed to promote both bonding and cooperation although only a limited number of studies have investigated this under natural conditions. Our aim was to assess the role of OT in bonding and cooperation in male Barbary macaques (*Macaca sylvanus*). First we tested for an effect of affiliation - grooming and triadic male-infant-male interactions - with bond and non-bond partners on urinary OT levels. Secondly we aimed to test whether grooming interactions (and thus increased OT levels) increase a male’s general propensity to cooperate in polyadic conflicts. We collected behavioral data via full-day focal animal protocols on 14 adult males and measured endogenous OT levels from 139 urine samples collected after affiliation and non-social control periods. Urinary OT levels were higher after grooming with any partner. By contrast, OT levels after male-infant-male interactions with any partner or with bond partners were not different from controls but were higher after interactions with non-bond partners. Previous grooming did not increase the likelihood of males to support others in conflicts. Collectively, our results support research indicating that OT is involved in the regulation of adult social bonds, including in non-reproductive contexts. However, our male-infant-male interaction results go against previous studies suggesting that it is affiliation with bond rather than non-bond partners that trigger the release of OT. Alternatively, OT levels may have been elevated prior to male-infant-male interactions thus facilitating interaction between non-bond partners. The lack of an association of grooming (and by extension increased OT levels) and subsequent support speaks against an OT linked increase in the general propensity to cooperate, yet further studies are needed for a more direct test including the possibility of partner-specific contingent cooperation.

## 1 Introduction

For mammals living in stable social groups, investing in strong social bonds can provide individuals with adaptive benefits (Ostner and Schülke, 2018) such as increased reproductive success (Cameron et al., 2009; Frère et al., 2010; Schülke et al., 2010; Strauss and Holekamp, 2019; Weidt et al., 2008) and increased survival (Archie et al., 2014; Giles et al., 2005; Silk et al., 2010). Social bonds are usually characterized by high rates of affiliative interactions (Ostner and Schülke, 2018) and may promote cooperative behavior (Schülke et al., 2010; Smith et al., 2011; Weidt et al., 2008; Young et al., 2014b) and buffer physiological impacts of stress (Cheney and Seyfarth, 2009; Young et al., 2014a).

One hormone implicated in the formation and maintenance of social bonds is the highly conserved neuropeptide oxytocin (OT). OT plays a role in promoting maternal behavior (Finkenwirth et al., 2016; Ross and Young, 2009) and ultimately a partner specific attachment between mother and offspring (Ross and Young, 2009). The oxytocinergic system is thought to have been co-opted from its ancestral function of mother-offspring attachment to also promote social bonds between adults (Numan and Young, 2016; Ziegler and Crockford, 2017). This has been best demonstrated in the context of pair bonds where OT helps regulate a social preference for a particular mating partner (French et al., 2018; Ross and Young, 2009). Oxytocin may also regulate social bonds more broadly outside the pair-bond and in non-reproductive contexts. In support of this, OT is released after positive, non-sexual, social interactions (chimpanzees, *Pan troglodytes*: Crockford et al., 2013; Preis et al., 2018; Wittig et al., 2014; tufted capuchins, *Sapajus apella*: Benítez et al., 2018; dogs, *Canis familiaris*: Romero et al., 2014). Because OT interacts with the reward system (Dölen et al., 2013; Love, 2014; Skuse and Gallagher, 2009), OT release potentially stimulates a ‘feel good’ sensation after positive social interactions. These sensations may be part of the mechanism by which social bonds are maintained (e.g., via emotional bookkeeping: Schino and Aureli, 2009).

OT release is partner specific, at least in some studies. In chimpanzees, urinary OT levels are increased after grooming with a bonded partner, but not after the same interaction with a non-bonded partner (Crockford et al., 2013). Similarly, in cooperatively breeding marmosets (*Callithrix jacchus*), strongly bonded family members show synchronous fluctuations in baseline urinary OT levels whereas weakly bonded partners do not, suggesting that affiliation with bond partners influences OT levels more than affiliation with non-bond partners (Finkenwirth et al., 2015). Other studies, however, suggest that OT is released independently of partner bond strength (Preis et al., 2018; Wittig et al., 2014) or that the impact of bond partner strength on OT secretion depends on the type of interaction (Wittig et al., 2014).

In addition to social bonding, oxytocin plays a key role in promoting coordination and cooperative behaviors under certain contexts. In economic games, intranasal administration of OT increases cooperation when participants had prior contact but not when they were anonymous (Declerck et al., 2010), and similarly in in-group but not out-group conditions (De Dreu et al., 2010; Ten Velden et al., 2017). Performance on a cooperative task was also improved by OT administration, suggestive of OT’s role in the facilitation of coordination of behavior (Arueti et al., 2013). These findings in humans are paralleled in chimpanzees where urinary OT levels were elevated during coordinated behaviors such as territorial border patrols (Samuni et al., 2017) and cooperative hunting (Samuni et al., 2018) compared to controls. Furthermore, the highest levels of urinary OT in chimpanzees occurred during inter-group encounters, which involve joint aggression against out-group members (Samuni et al., 2017). Depending on context, OT appears to facilitate participation in polyadic aggression through increased coordination and in-group favoritism in humans and chimpanzees. The generality of these patterns beyond these taxa remains to be tested.

We aimed to investigate the role of oxytocin in the maintenance of social bonds and cooperation in male macaques. Macaque males of several species form strong, equitable and stable social bonds with other males (Kalbitz et al., 2016; Young et al., 2014b) which are predictive of cooperation via coalition formation (Berghänel et al., 2011; Schülke et al., 2010; Young et al., 2014b). Coalitions serve to increase or maintain male dominance rank (Young et al., 2014c), and increase mating success (Küster and Paul, 1992; Young et al., 2013) or reproductive success (Schülke et al., 2010). In addition to male-male bonds, macaques may also from strong male-female bonds (Haunhorst et al., 2016; Massen and Sterck, 2013). Males frequently support females in agonistic conflicts (Haunhorst et al., 2017; Kulik et al., 2012; Small, 1990), and similarly to male-male relations, the probability to support is predicted by social bond strength (Haunhorst et al., 2017; Kulik et al., 2012).

A behavioral pattern proposed to enhance male social bonding are triadic male-infant-male interactions which are characteristic yet not exclusive to Barbary macaques (Deag, 1980; Paul et al., 1996). These interactions are ritualistic in nature and involve two males sitting in body contact holding an infant in between them while teeth-chattering and often inspecting the infant’s genitals (Deag, 1980; Deag and Crook, 1971). This behavior most commonly involves newborn infants, although yearlings and two-year-olds are also sometimes involved (Paul et al., 1996). Triadic male-infant-male interactions have been proposed to be used as a tool to enhance male-male social bonds (Henkel et al., 2010; Kalbitz et al., 2017; Paul et al., 1996) and predict coalition formation in the mating season (Berghänel et al., 2011). Alternatively, though not mutually exclusive, male-infant-male interactions may be used as a form of ‘agonistic buffering’ (Deag, 1980; Deag and Crook, 1971; Paul et al., 1996). Thus, triadic male-infant-male interaction is a behavior with qualities similar to others that induce the release of OT.

Similar to previous studies in chimpanzees and tufted capuchins (Benítez et al., 2018; Crockford et al., 2013), we tested in Barbary macaques whether urinary OT levels were influenced by grooming interactions and - given its functional relevance in Barbary macaques - also by male-infant-male interactions. We predicted that urinary OT levels would be higher after affiliative interactions (i.e., grooming, male-infant-male interactions) compared to a control period without any social interactions. We additionally tested whether the release of OT was partner-specific (i.e., bond partners vs. non-bond partners). If OT release is partner-specific, we predicted that it will be higher after affiliations with bond partners than with non-bond partners. A secondary aim was to test whether OT would generally increase the propensity to cooperate in within-group polyadic agonistic conflicts. To do this under natural conditions, we first determined which affiliative behaviors are positively related to urinary OT levels (as part of the first aim), to use the occurrence of this interaction as a proxy for elevated OT levels in the subject. We consequently predicted, that the probability to cooperate, i.e. accept a solicitation to support another individual in an agonistic conflict, would be higher after an affiliative interaction.

## 2 Materials and Methods

### 2.1 Study site and animals

Study subjects belonged to one of three groups of Barbary macaques living together in 14.5 ha. of enclosed forest at Affenberg Salem, Germany (de Turckheim and Merz, 1984). Monkeys were provisioned once daily with fruits, vegetables, grains and had *ad libitum* access to water and monkey chow. Data collection took place from 31 March to 17 December 2016, including one non-mating season (31 March to 26 October) and one mating season (27 October to 17 December). The start of the mating season was defined by the first observed ejaculatory copulation. The study group (group C) consisted of 13-14 adult males (one male died during the study period), 20 adult females, 2 large subadult males, 8 immature males, 10 immature females and 1 newborn infant male. All members of the group were individually recognized by observers.

### 2.2 Behavioral data collection

Behavioral data were collected from 14 adult males using continuous focal animal sampling (Martin and Bateson, 2007) during individual full-day focal animal follows, in which the occurrence and partners of all social interactions were recorded (Total = 4355 hours, 311 ± SD 40 hours per individual).

### 2.3 Assessing dyadic bond strength

To assess dyadic bond strength, we calculated the dyadic Composite Sociality Index (CSI; Silk et al., 2010, 2006), with slight modifications as described in Haunhorst et al. (2016). This index ranges from 0 to infinity and has a mean value of 1, where higher CSI scores indicate a stronger social bond. To calculate the CSI, we chose seven significantly correlated affiliative behavioral variables: duration and count of close proximity (≤ 1.5 meters) without aggression, duration and count of body contact, duration and count of grooming and count of triadic male-infant-male interactions. Both duration and count of behaviors were corrected for the total observation time of the dyad. Male Barbary macaques affiliate with females much more frequently than with other males (mean ± SD behavior seconds per observation hour per sex of dyad: proximity, male-male dyads = 69 ± 57; proximity, male-female dyads = 372 ± 121; body contact, male-male dyads = 36 ± 38; body contact, male-female dyads = 247 ± 71; grooming, male-male dyads = 22 ± 20; grooming, male-female dyads = 201 ± 55). If we had included both male-male and male-female dyads into a single CSI scores this would lower the CSI scores for male-male dyads. As a results we would potentially miss-classify some male-male dyads as non-bond when in fact they are bond partners, given that male-male bonds are meaningful and have adaptive significance (see introduction). Therefore, we constructed separate CSI scores for male-male and male-female dyads. The two large subadult males in our study group were included in the calculation of the male-male CSI scores because they supported other adult males in agonistic conflicts. Furthermore, we calculated separate CSI scores for the non-mating and mating seasons as affiliation patterns may change across seasons. Out of the seven affiliative behavior conditions, we only included them in the CSI calculation if their mean frequency of occurrence per dyad in each period was > 2 to avoid rare behaviors disproportionately affecting the CSI scores. We defined bond partners as those dyads with a CSI score > 1 (above the group mean).

### 2.4 Urine sample collection

Urine samples were collected opportunistically from individuals during focal follows. When monkeys were seen to urinate, the urine was caught with a plastic bag when possible or collected from leaves, branches, rocks or the ground by using a disposable pipette or salivette (Salivette Cortisol, Sarstedt, Nümbrecht, Germany). The use of salivettes to collect urine has recently been validated and successfully applied to urine samples from free-ranging macaques (Danish et al., 2015; Müller et al., 2017). Urine samples contaminated with feces, blood or urine from other individuals were not collected. Urine samples collected by pipette were transferred to 2 ml cryotubes. Both samples stored in cryotubes and salivettes were kept in a thermos flask filled with ice while in the field. At the end of the day, urine was recovered from the salivettes by centrifugation for 5 min at 1500 rpm using an electric centrifuge and also transferred to 2 ml cryotubes. Samples were split into two aliquots (100 to 2000 μl each). One aliquot was used for analysis of creatinine. In the second aliquot, 0.5 N phosphoric acid were added to urine at a ratio of 1:10 acid to urine to prevent the breakdown of OT in the sample (Reyes et al., 2014; Ziegler, 2018). All samples were then stored in a freezer at −20°C. When data collection was complete, samples were transported in containers with dry ice to the lab and stored once again at −20°C.

Urine samples were collected from all 14 adult males (7 to 25 years old) of the study group. We presumed a clearance window of 15 to 60 min for excretion of OT in urine, as done in previous studies investigating urinary OT levels in other non-human primates which show biologically relevant changes in behavior during this window (Benítez et al., 2018; Crockford et al., 2013; Samuni et al., 2017). Studies in humans and marmosets have demonstrated elevated OT levels in urine 30 to 60 after administration of radio-labelled hormone (humans: Amico et al., 1987; marmosets: Seltzer and Ziegler, 2007). Exogenous administration of OT in tufted capuchin monkeys also caused elevated urinary OT levels 15-60 min after administration (Benítez et al., 2018). Prior to analysis, urine samples were assigned to different behavioral conditions depending on whether at least one grooming (total time ≥ 60 sec), triadic male-infant-male interaction or no social interactions occurred in the 45 min clearance window. As we were interested in the role of OT in bonding in a non-sexual context, we only considered samples collected during the non-mating season for analysis. Furthermore, samples were excluded from analysis if any ejaculatory copulations, play or coalitions co-occurred in the clearance window because these behaviors could potentially influence OT levels and confound results. This left us with 76 non-social (control) samples (mean = 5.8, range = 2-11 per individual) and 63 samples where at least one affiliation occurred (test samples: mean = 4.8, range = 1-9 per individual).

### 2.5 Extraction and hormone analysis

The extraction and analysis of OT followed a protocol described in detail in Samuni et al. (2017). Briefly, urine samples were thawed and kept cool using an Iso-rack (0°C; Eppendorf). Then samples were centrifuged for 1 min at 1500 rpm at 4°C. Solid-phase extraction cartridges (Chromabond HR-X, 30mg, 1 ml, Macherey-Nagel, Dueren, Germany) were conditioned with 1 ml MeOH followed by 1 ml distilled HPLC-water. Cartridges were then filled with up to 1 ml dilution buffer (water, 0.1% TFA) and 20 to 100 μl of urine. Diluted urine was allowed to run through the cartridge. Then, the cartridge was washed with 1 ml washing solution (10% ACN, 1% TFA) and dried using a vacuum. Hormones were eluted using 1 ml ACN 80% into clean test tubes. Elutes were evaporated at 50°C with pressurized air. Then 300 μl EtOH 100% was added to each test tube and shaken gently. Test tubes were allowed to sit for 1 hour at 4°C to precipitate proteins before being evaporated again at 50°C. Samples were then reconstituted with 250 μl assay buffer from a commercially available enzyme immunoassay kit (Assay Designs; 901-153A-0001), and vortexed gently for 10 sec by hand. Extracts were then transferred to 1.5 ml labeled eppendorf tubes, and vortexed for 1 min at 10,000 rpm. Extracts were then kept cool on ice while preparing the assay. The assay was then performed according to instructions provided by the manufacturer.

To determine the efficiency of the extraction protocol, we created 5 pools of Barbary macaque urine samples. Before extraction, 75 μl of each pooled sample were spiked with 75 μl of an OT standard (1500 pg/ml). We used the values from the spiked and unspiked samples to calculate percent recovery for extraction efficiency and assay accuracy following the formula given in Behringer et al. (2012). Mean extraction was 81.0% (range: 92.7%-68.7%, SD = 10.2, N = 5). We investigated matrix effects that could potentially interfere with the assay system by testing for parallelism. Out of a pool sample, we took 3 ml of urine and extracted them according to our extraction protocol. Of the resulting 500 μl of extract, 250 μl were taken and serially diluted. Another 1 ml of the urine pool sample was mixed with 100 μl of an OT standard solution (10 000 pg/ml), extracted and serially diluted as described above. Dilutions of the spiked and unspiked pool sample were then brought to assay. Serially diluted pool samples of spiked and unspiked Barbary macaque urine were parallel to the standard as confirmed by visual inspection (Fig. S1).

The assay standard curve ranged from 15.62 to 1000 pg/ml and assay sensitivity at 90% binding was 30 pg/ml. Intra-assay coefficients of variation (CV) of high and low value quality controls were 5.2% (high) and 31.3% (low) while respective figures for inter-assay CVs were 11.0% (high) and 19.7% (low).

Urinary OT concentrations were corrected for levels of creatinine to account for differences in volume and concentration of excreted urine (Bahr et al., 2000), and are expressed as pg/mg creatinine. Because very low concentrations of creatinine may lead to an overestimation of hormone concentration we excluded all samples (N = 3) with < 0.5 mg/ml creatinine.

### 2.6 Statistical analysis

To test whether affiliative interactions influenced urinary OT levels, we fitted two Bayesian multilevel linear regression models (model 1a, b) with a Gaussian response distribution and identity link function. To test whether the probability to give support in an agonistic encounter after being recruited was influenced by a previous grooming interaction, we fitted a Bayesian multilevel linear regression model with a Bernoulli response distribution and logit link function. We included male identity as a random effect in all models. In all models, predictor variables varied within male identity and therefore we included random slopes as well as correlation parameters between random intercepts and random slopes into the models (Barr et al., 2013; Schielzeth and Forstmeier, 2009). We fitted models using the computational framework Stan (https://mc-stan.org), called via R (version 3.5.2; R Core Team, 2018) by using the function brm from the package brms (version 2.9.0; Bürkner, 2017). We ran all models with 5000 iterations over four MCMC chains including an initial 1000 “warm up” iterations for each chain, resulting in a total of 16000 posterior samples (Bürkner, 2017). In all models, we deemed the MCMC results as reliable because there were no divergent transitions during warm up, all Rhat values were equal to 1.00 and visual inspection of a plot of the chains showed that they were able to converge. We used a set of weakly informative priors to improve convergence, guard against overfitting and regularize parameter estimates (Lemoine, 2019; McElreath, 2016): for the intercept and beta coefficients we used a normal distribution with mean 0 and standard deviation 10; for the standard deviation of group level (random) effects and sigma we used a Half-Cauchy distribution with location 0 and scale parameter 1; for the correlation between random slopes we used LKJ Cholesky prior with eta 2.

For all models, we report the estimate as the mean of the posterior distribution and 95% credible intervals (CI). We calculated the proportion of the posterior samples that fall on the same side of 0 as the mean. This may be interpreted as the probability (Pr) that a given predictor was associated with an outcome, where Pr = 1 indicates that the estimate was entirely positive or negative and Pr = 0.5 indicates that the estimate is centered around 0 and thus the predictor likely had no effect.

#### 2.6.1 Effect of affiliation on urinary OT levels

To test whether urinary OT levels were generally influenced by affiliative behaviors we fitted two models. As the response we log-transformed urinary OT levels to achieve a more symmetrical distribution. In model 1a, we tested for a general effect of affiliation and included one categorical predictor where OT levels following grooming and triadic male-infant-male interactions were compared to non-social controls. In model 1b, we tested whether OT levels would be influenced differently after affiliation with bond versus non-bond partners. Therefore, we split samples after triadic male-infant-male interactions into bond and non-bond partner categories. As we were only able to collect two urine samples where focal males groomed with a non-bond partner, we decided not to split grooming samples according to partner bond strength.

#### 2.6.2 Effect of grooming on probability to give support

To test whether the probability to give support in an agonistic encounter (between adult and/or subadult individuals) after being recruited was influenced by a previous grooming interaction, we fitted one model. As the response, we included whether our focal male supported another adult individual following a recruit attempt (no/yes). As a test predictor, we included whether our focal animal was in a grooming interaction (≥ 60 sec) with an adult individual within 15-60 min before the recruit behavior (no/yes). This time window was chosen because intranasal administration of OT in rhesus macaques influenced social behaviors up to two hours after inhalation (Chang et al., 2012). Therefore, we chose a comparatively conservative window of 15-60 min for when naturally centrally released OT may still exert behavioral effects. As a control predictor, we included the bond strength of the focal animal to the recruiter (non-bond/bond).

## 3 Results

We first tested for a general effect of affiliative interactions (grooming and male-infant-male triadic interactions) irrespective of partner bond strength on urinary OT levels. Urinary OT levels were substantially higher after grooming with any partner compared to non-social controls (mean ± SD OT: non-social: 357 ± 400 pg/mg creatinine; grooming: 589 ± 612 pg/mg creatinine; average increase of 65%; Pr = 0.97; Table 1 a, b; Fig. 1; Fig. 2), while this was not the case for male-infant-male interactions with any partner (mean ± SD OT: male-infant-male: 455 ± 438 pg/mg creatinine; Pr = 0.77; Table 1).

**Table 1:** Results of models 1a and 1b testing effect of different affiliation conditions on urinary OT levels. In both models, male identity was included as a random effect, N = 13 males, N = 139 samples. CI = 95% credible intervals, Pr = proportion of the posterior samples that fall on the same side of 0 as the mean.

|  | Estimate | SD | CI lower | CI upper | Pr |
| --- | --- | --- | --- | --- | --- |
| **(a)** |  |  |  |  |  |
| Intercept | 5.48 | 0.12 | 5.25 | 5.71 | 1.00 |
| Groom | 0.47 | 0.25 | -0.01 | 0.99 | 0.97 |
| Male-infant-male | 0.17 | 0.23 | -0.30 | 0.60 | 0.77 |
| **(b)** |  |  |  |  |  |
| Intercept | 5.49 | 0.11 | 5.26 | 5.70 | 1.00 |
| Groom | 0.47 | 0.25 | -0.01 | 1.00 | 0.97 |
| Male-infant-male bond | -0.05 | 0.28 | -0.59 | 0.51 | 0.58 |
| Male-infant-male non-bond | 0.57 | 0.33 | -0.09 | 1.22 | 0.96 |

**Fig. 1:**
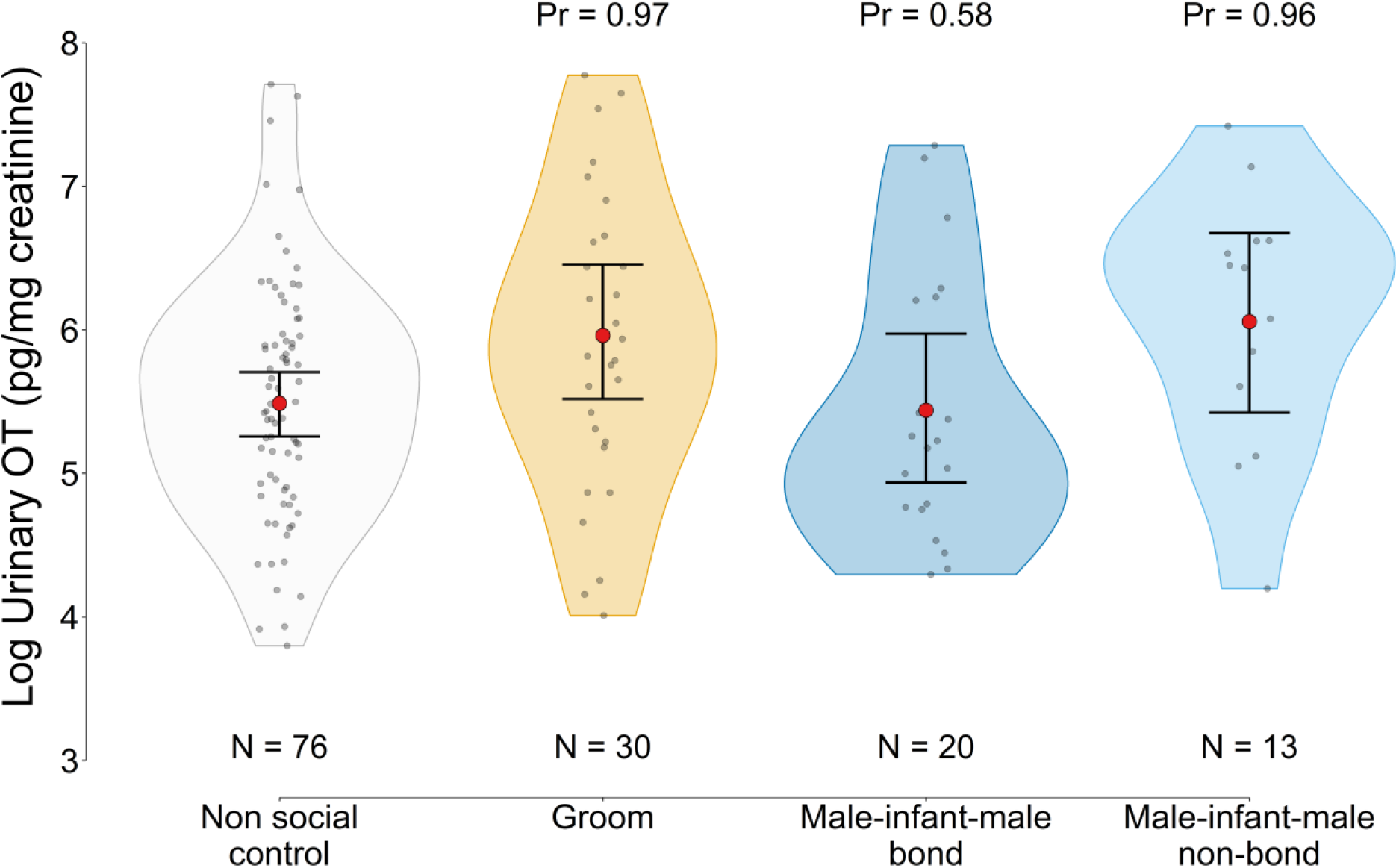
Urinary OT levels per behavioral condition. Violin plots show the density of observed data points. Solid red dots show fitted values from model 1b: mean of posterior distribution and 95% credible intervals. Pr = proportion of the posterior samples that fall on the same side of 0 as the mean. N = number of samples per condition.

**Fig. 2:**
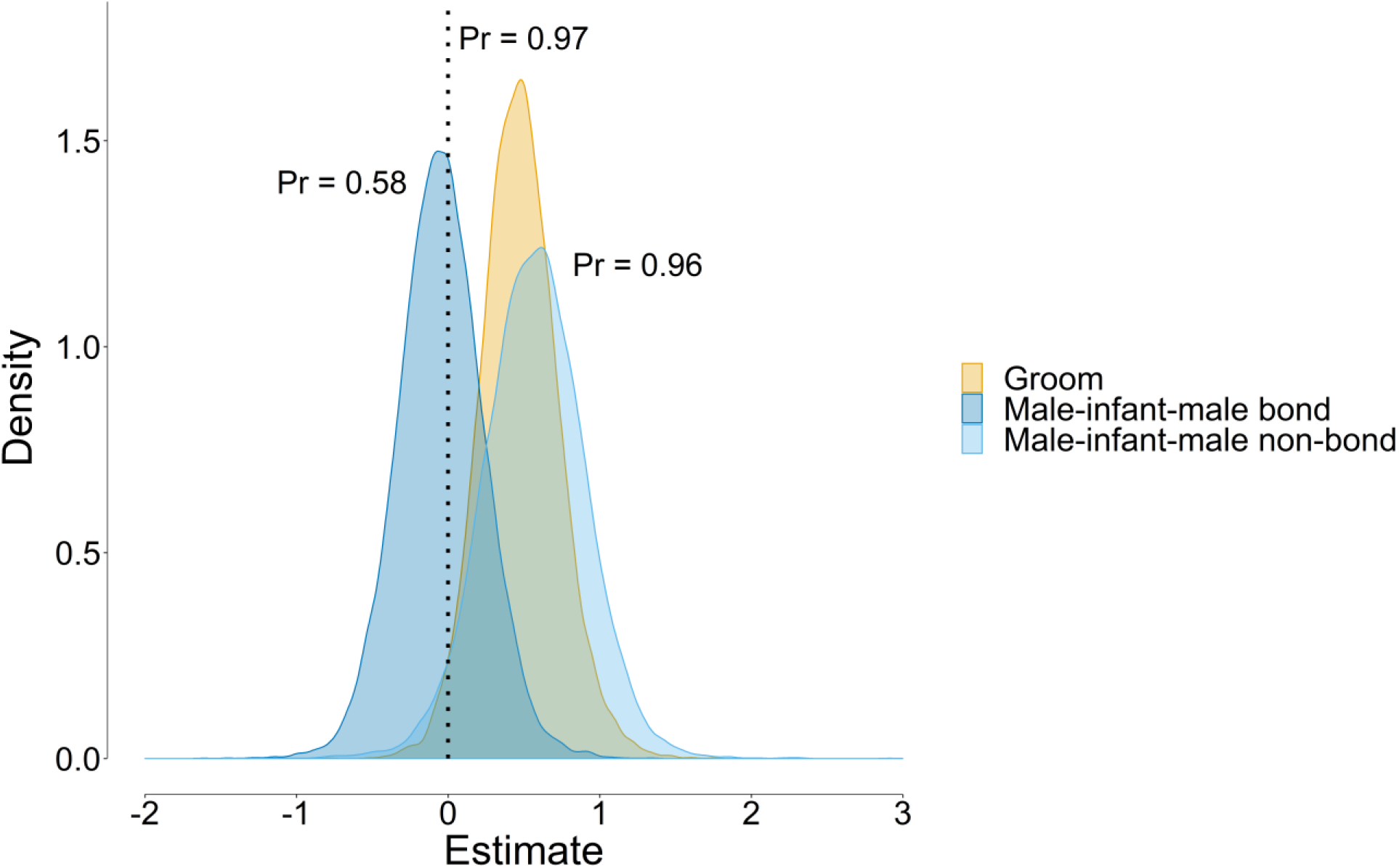
Posterior probability distribution of the difference in urinary OT levels after different affiliative behaviors compared to non-social controls. Pr = proportion of the posterior samples that fall on the same side of 0 as the mean.

When we separated male-infant-male interaction samples by bond strength, urinary OT levels after male-infant-male interactions with bond partners were also not substantially different from non-social controls (mean ± SD OT: non-social: 357 ± 400 pg/mg creatinine; male-infant-male bond: 360 ± 410 pg/mg creatinine;). In contrast, urinary OT levels were substantially higher after triadic male-infant-male interactions with non-bond partners than non-social controls (mean ± SD OT: male-infant-male non-bond: 600 ± 456 pg/mg creatinine; average increase of 68%; Pr = 0.96; Table 1 b; Fig. 1; Fig. 2).

### 3.1 Effect of grooming on probability to give support

We recorded a total 205 attempts of adult individuals to recruit the focal animal for an agonistic conflict. In 64 (31%) cases the focal animal supported the recruiter and in 67 (33%) cases the focal animal was in a grooming interaction with any adult group member 15 to 60 minutes prior to the recruitment attempt (these samples are not mutually exclusive). In only 7 (3%) cases were the previous grooming partner also the recruiter. Grooming interactions did not substantially influence the probability to support a recruiter in an agonistic encounter within 15 to 60 minutes after the grooming interaction (N support given when groomed before = 16, N support given when not groomed before = 48; Pr = 0.86; Table 2).

**Table 2:** Model 2 results testing the effect of grooming on the probability to give support in an agonistic conflict after being recruited. Bond strength with the recruiter was included as a control variable. N = 14 males, N = 205 observations. CI = 95% credible intervals, Pr = proportion of the posterior samples that fall on the same side of 0 as the mean.

|  | Estimate | SD | CI lower | CI upper | Pr |
| --- | --- | --- | --- | --- | --- |
| Intercept | -1.05 | 0.35 | -1.80 | -0.41 | 1.00 |
| Groom before? (yes) | -0.45 | 0.43 | -1.32 | 0.37 | 0.86 |
| Recruiter bond (bond) | 0.81 | 0.46 | -0.03 | 1.82 | 0.97 |

## 4 Discussion

Overall we found a high probability that urinary OT levels are elevated following grooming interactions in adult male Barbary macaques. This is generally in line with previous studies showing a positive relationship between OT and grooming (primates: Benítez et al., 2018; Crockford et al., 2013; Snowdon et al., 2010; vampire bats, *Desmodus rotundus*: Carter and Wilkinson, 2015), as well as other socio-positive interactions more generally (primates: Preis et al., 2018; Benítez et al., 2018; Snowdon et al., 2010; Wittig et al., 2014; vampire bats: Carter and Wilkinson, 2015; dogs: Romero et al., 2014). Given the low number of grooming between non-bonded partners in our study, we could not test partner specific effects of grooming. In chimpanzees, oxytocin release was partner-specific in one population, with elevations after grooming with a bond, yet not with a non-bond partner (Crockford et al., 2013); in another population OT levels were generally increased after affiliation (including grooming) irrespective of partner bond strength (Preis et al., 2018). Relationship quality was tested differently in these two studies, with relationship quality being either categorized dichotomously into bond and non-bond partners (Crockford et al., 2013), or being tested on a continuous scale (Preis et al., 2018). Our cut-off relationship strength value for classification as a bond partner was much lower than the one used for chimpanzees. We do not know how nonhuman primates classify each other into biologically meaningful bond and non-bond categories, e.g. an inner clique of 2-3 bonded partners (Hill et al., 2008; Zhou et al., 2005), and if this mental classification mediates OT release. In principle there is good evidence that classification into bond partners affects physiological responses to social interactions. The social buffering phenomenon shows that the presence or interaction with closely bonded partners during stressful events mitigates the release of glucocorticoids (Hennessy et al., 2009; Kikusui et al., 2006; Wittig et al., 2016; Young et al., 2014a) with OT release mediating social buffering of the stress response (Crockford et al., 2017; Hennessy et al., 2009; Kikusui et al., 2006; Smith and Wang, 2014).

Unexpectedly, urinary OT levels were elevated after triadic male-infant-male interactions with non-bond partners, but not after interactions with bond partners. This finding contradicts the idea that it is affiliation with bond rather than non-bond partners that triggers the release of OT (Crockford et al., 2013; Finkenwirth et al., 2015). This could indicate that male-infant-male interactions serve to promote the formation of social bonds with not yet bonded partners, while physiologically not impacting interactions between established partners. An untested, yet possible alternative given the correlational nature of our study is the reversed cause-effect directionality: instead of a male-infant-male interaction triggering the release of OT, OT may increase the probability of a male-infant-male interaction to occur. In this scenario, male-infant-male interactions do not function in bond formation, but for other reasons, for example as a form of “agonistic buffering” (Deag, 1980; Deag and Crook, 1971; Paul et al., 1996). In support of this idea, rates of male-infant-male interactions increase during tense feeding situations while other types of affiliation (such as grooming) decrease (Paul et al., 1996). Social relationships between adult males are generally tense and affiliation between them often takes place in the presence of infants (Deag, 1980; Preuschoft and Paul, 2000). Due to the anxiolytic effect of OT (Neumann and Landgraf, 2012), elevated levels of OT may facilitate male-infant-male interactions via increasing the motivation to approach and at the same time reducing avoidance behaviors toward other males (Kemp and Guastella, 2011). Reduced anxiety may be particularly useful for interacting with non-bond males with whom the relationship is presumably more tense and unpredictable than with bond partners (Young et al., 2014b). Such an explanation would be consistent with our finding that OT levels were elevated after male-infant-male interactions with non-bond partners. While intriguing, we need to stress our small sample size of male-infant-male interactions with non-bond partners as well as previous work on this and other species pointing to the bond strengthening and cooperation-enhancing function of male-infant-male interactions (Berghänel et al., 2011; Kalbitz et al., 2017). Thus, additional studies able to disentangle OT levels directly before and after affiliative interactions are clearly needed for a more conclusive picture.

Prior grooming did not increase the probability of supporting a group member in a conflict. This test builds on the assumption that after engaging in grooming OT levels will be elevated, potentially influencing behavior and more specifically cooperative tendency. If true, this finding suggests that OT did not increase a male’s general tendency to cooperate in a conflict, which is maybe not surprising given the wealth of studies indicating that OT’s prosocial effects depend on situational context and interaction partner (Bartz et al., 2011). In an economic game, human cooperation was enhanced by intranasal OT administration only if participants had prior contact yet not with strangers (Declerck et al., 2010), and similarly, female house mice (*Mus musculus domesticus*) receiving OT actually decreased the propensity to cooperate in communal breeding with strangers (Harrison et al., 2017). In another study in humans intranasal OT increased trust but not if the partner was portrayed as untrustworthy (Mikolajczak et al., 2010). Thus, the social information on a partner is an important component for OT-induced cooperation and depending on this information OT may reduce the propensity to cooperate.

Our results do not exclude the possibility that OT promotes direct or partner-specific cooperation. From a behavioral perspective, individuals are more likely to support others with whom they have groomed in the recent past (long-tailed macaques, *Macaca fascicularis*: Hemelrijk, 1994; chacma baboons, *Papio ursinus*: Cheney et al., 2010). In our study there were only a few cases of the former grooming partner asking for help within the next hour, therefore we could not explicitly test this scenario. Contingent cooperation appears to be rare in animals, and more commonly support is given less strictly on a contingent basis but instead to bonded partners who form long term alliances (Cheney, 2011). OT would then mediate cooperation with specific partners through its role in promoting the formation of social bonds. While we did not have an explicit aim to test the effect of partner bond strength on the probability to cooperate, this variable was included in our model as a control predictor, and it did substantially increase the probability of giving support. Preferentially giving support to bonded partners has also previously been shown in this (Young et al., 2014b) and other species (Schülke et al., 2010; Smith et al., 2011; Watts, 2002).

Overall, our study adds to the body of research indicating that OT is involved in the regulation of adult social bonds, including in non-reproductive contexts. Questions still remain under which contexts OT release is partner specific. It has been suggested that in smaller social groups, all group members are bonded to a sufficient degree to elicit OT release after affiliative interactions (Benítez et al., 2018), whereas in larger groups variance in affiliation rates may be large enough that OT release may only occur after affiliations with more closely bonded partners. Such an explanation would be consistent with observed differences in partner specificity of OT release between chimpanzee populations (Crockford et al., 2013; Preis et al., 2018). Finally, the lack of an effect of male-infant-male interactions on OT levels in male Barbary macaques, at least with bond partners and perhaps overall, remains puzzling. Particularly so given that behaviors other than grooming that potentially promote social bonding also increased urinary OT levels in other species (Benítez et al., 2018; Romero et al., 2014; Wittig et al., 2014). One difference between grooming and male-infant-male interactions is that male-infant-male interactions are more ritualistic in nature. Perhaps the bonding effect of ritualized behavior may be under the control of other neuropeptides, such as endorphins, as has been shown for synchronous dancing in humans (Tarr et al., 2015, 2016).

## Acknowledgements

We thank Ellen Merz and Roland Hilgartner for permission to conduct the study at Affenberg Salem. We thank Lauren Cassidy, Tatjana Kaufmann and Lilah Sciaky for help with urine sample and behavioral data collection. We are grateful to Vera Schmeling for general assistance in the lab. This research was funded by the Deutsche Forschungsgemeinschaft (DFG, German Research Foundation) - Project number 254142454 / GRK 2070.

