## Supplementary figures and images for "Oxytocin increases after affiliative interactions in male Barbary macaques"

### Supplemental Figure 1

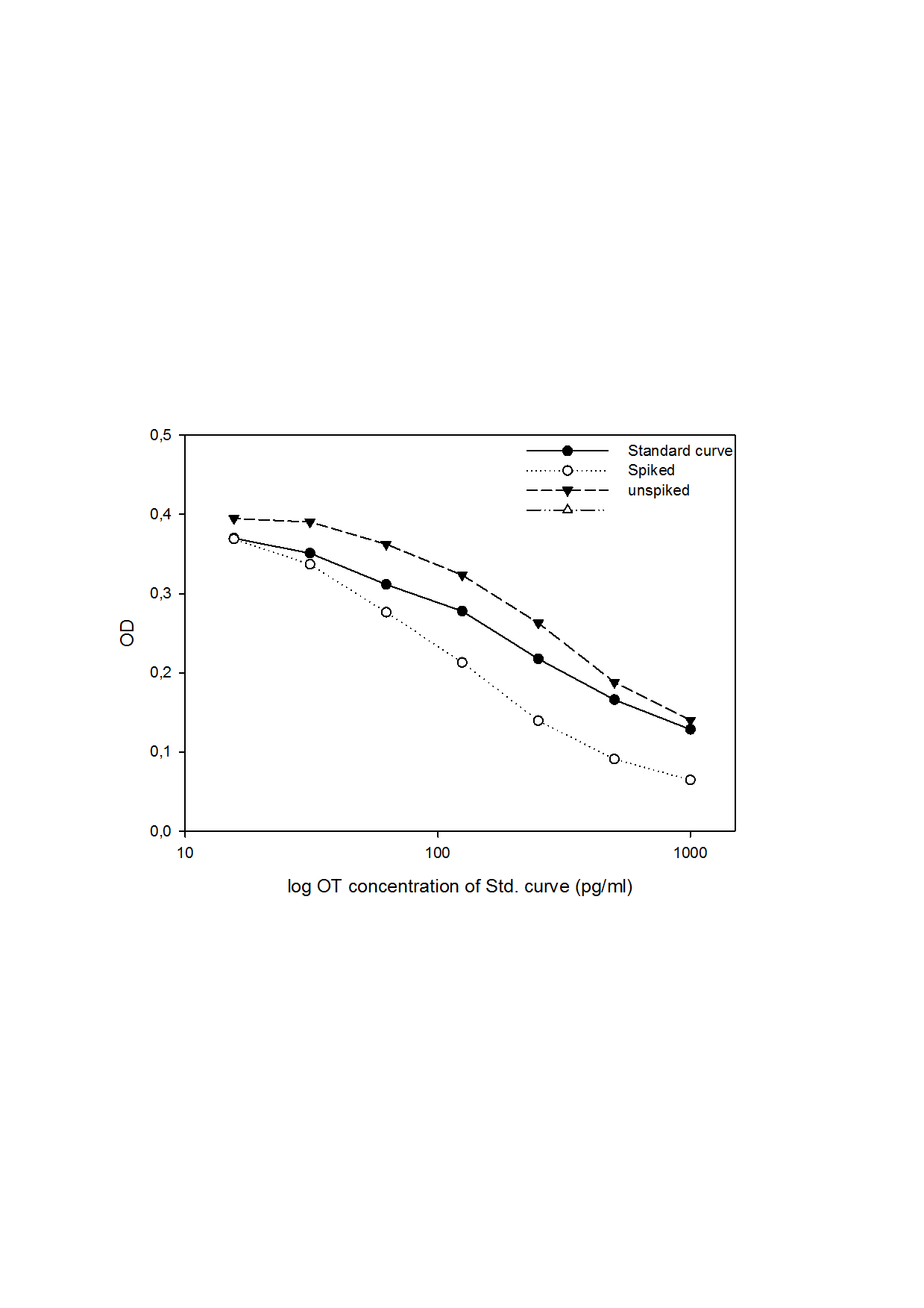
